# Characterization of the evolutionary dynamics of influenza A H3N2 hemagglutinin

**DOI:** 10.1101/2020.06.16.155994

**Authors:** Maggie Haitian Wang, Jingzhi Lou, Lirong Cao, Shi Zhao, Paul KS Chan, Martin Chi-Wai Chan, Marc KC Chong, William KK Wu, Renee WY Chan, Yuchen Wei, Haoyang Zhang, Benny CY Zee, Eng-kiong Yeoh

## Abstract

Virus evolution drives the annual influenza epidemics in human population worldwide. However, it has been challenging to evaluate the mutation effect of the influenza virus on evading the population immunity. In this study, we introduce a novel statistical and computational approach to measure the dynamic molecular determinants underlying epidemics by the effective mutations (EMs), and account for the time of waning mutation advantage against herd immunity by the effective mutation periods (EMPs). Extensive analysis is performed on the genome and epidemiology data of 13-year worldwide H3N2 epidemics involving nine regions in four continents. We showed that the identified EM processed similar profile in geographically adjacent regions, while only 40% are common to Europe, North America, Asia and Oceania, indicating that the regional specific mutations also contributed significantly to the global H3N2 epidemics. The mutation dynamics calibrated that around 90% of the common EMs underlying global epidemics were originated from South East Asia, led by Thailand and India, and the rest were originated from North America. New Zealand was found to be the dominate sink region of H3N2 circulation, followed by UK. All regions might act as the intersection in the H3N2 transmission network. The proposed methodology provided a way to characterize key amino acids from the genetic epidemiology point of view. This approach is not restricted by the genomic region or type of the virus, and will find broad applications in identifying therapeutic targets for combating infectious diseases.

## Introduction

Rapid mutations in the influenza virus have led to annual influenza epidemics worldwide, affecting 5–15% of the global population^1,2^. Notwithstanding increasing evidence regarding the evolution of the influenza virus, some important questions remain unanswered, particularly those regarding the quantification of key mutations underlying local epidemics and the persistence of a virus mutation for evasion from herd immunity. Responses to these questions would help understand evolutionary characteristics of influenza viruses and the factors influencing their activities. Studies on the evolution of the influenza virus have focused on genetic or antigenic aspects. At the genetic level, on comparing synonymous and non-synonymous mutations, codons under positive selection pressure have been identified^3,4^; analysis of the genetic distances among viral genomes via phylogenetic analysis have revealed relationship among viral strains^5,6^. At the antigenic level, through the standard hemagglutination inhibition (HI) assay, specific antibody titers against a particular influenza strain were detected in biological samples^7^; and transformation of assay reads on the antigenic map facilitated the detection of antigenic clusters of the viral strains^8,9^. However, the disease activity of influenza at the population level, which plays a critical role in the selection of evolutionarily advanced genetic strains, has not been systematically incorporated in the evaluation of influenza virus substitutions.

This study proposes a novel statistical and computational framework to evaluate the important individual amino acid substitutions in influenza epidemics. In the proposed framework, effective mutations (EMs) are defined as the amino acid substitutions significantly associated with the prevalence of a viral serotype in the population and the effective mutation period (EMP) is defined as the corresponding period of contribution of an EM. Therefore, EMs are different from positively selected codons identified solely on the basis of genetic variations or the canonical antigenic sites that have been structurally and experimentally localized^10,11^. The EMs are not subject to HI readings and are specific to epidemics by time and region. We applied this framework on human influenza A/H3N2 virus harvested between 2003 and 2018. These data comprise 9,827 haemagglutinin (HA) sequences and seropositivity rates were determined from 4.1 million specimens. Nine regions on four continents were investigated, including New York (NY), California (CA), United Kingdom (UK), provinces in South China (SCP), Hong Kong SAR (HK), Singapore (SG), India (IN), Thailand (TH) and New Zealand (NZ). Distinct features of viral evolution dynamics were observed among continents. The EM and EMP calibrated mutation tempo at amino acids level, and refined the global transmission pathways of the H3N2 influenza virus, particularly in the South East Asia (SEA) area.

## Results

### Effective Mutations (EMs) – key substitutions underlying the epidemics

First, we used data from California (CA) as an example to study the mutation characteristics of a region and compared the EMs identified in different parts of the world. In CA, from influenza season 2005/06 to 2017/18, two to three EMs on average underlay annual A/H3N2 epidemics (**Figure 1a**). The punctuated antigenic drift of the A/H3N2 virus was reflected in the scale of EMs, i.e., more EMs were observed during large epidemic waves. For example, during the peak of H3N2 in 2012/13, a set of 12 amino acid substitutions was identified (**Figure 1a, blue box**), wherein EMs (49, 61, 64, 161, 214, 239, 294), including those in codon 214 and 294, which are known antigenic sites, were detected in all nine regions (**Figure 1, red arrows**). This shows that the proposed method can consistently identify the important amino acids in independent data sets. Furthermore, the subset (19, 158, 505) was also synchronized and repeatedly captured in NY, UK, SCP, NZ and IN (**Figure 1, blue arrows**). Among these EMs, codon 158 is one of the seven experimentally validated amino acids proximal to receptor-binding sites and causes antigenic variations in the H3N2 virus^12^, and codon 505 was located in the stem region of the HA protein. Antigenic sites on the conserved regions in the viral genome were particularly difficult to detect via the HI assay, as their effects are usually masked by the immune dominance of the HA1 region. While we evaluated substitutions in set-form, novel epistatic loci contributing to overall viral fitness in population epidemics might be detected. Furthermore, codon 241 is a potential new antigenic site identified by this framework. Though not previously reported, codon 241 was identified as an EM in all nine regions; it mutated twice in NY and CA during the study period (**Figure 1a-b**), and is located at the outer loop on the globular head of the HA protein. The results suggest that this genetic epidemiology framework might be used to efficiently screen for novel loci underlying infectious disease epidemics. The identified amino acids provide a list of immune-relevant targets for vaccine and therapeutics development. A complete list of EMs is provided in the Table S1.

**Figure 1.**
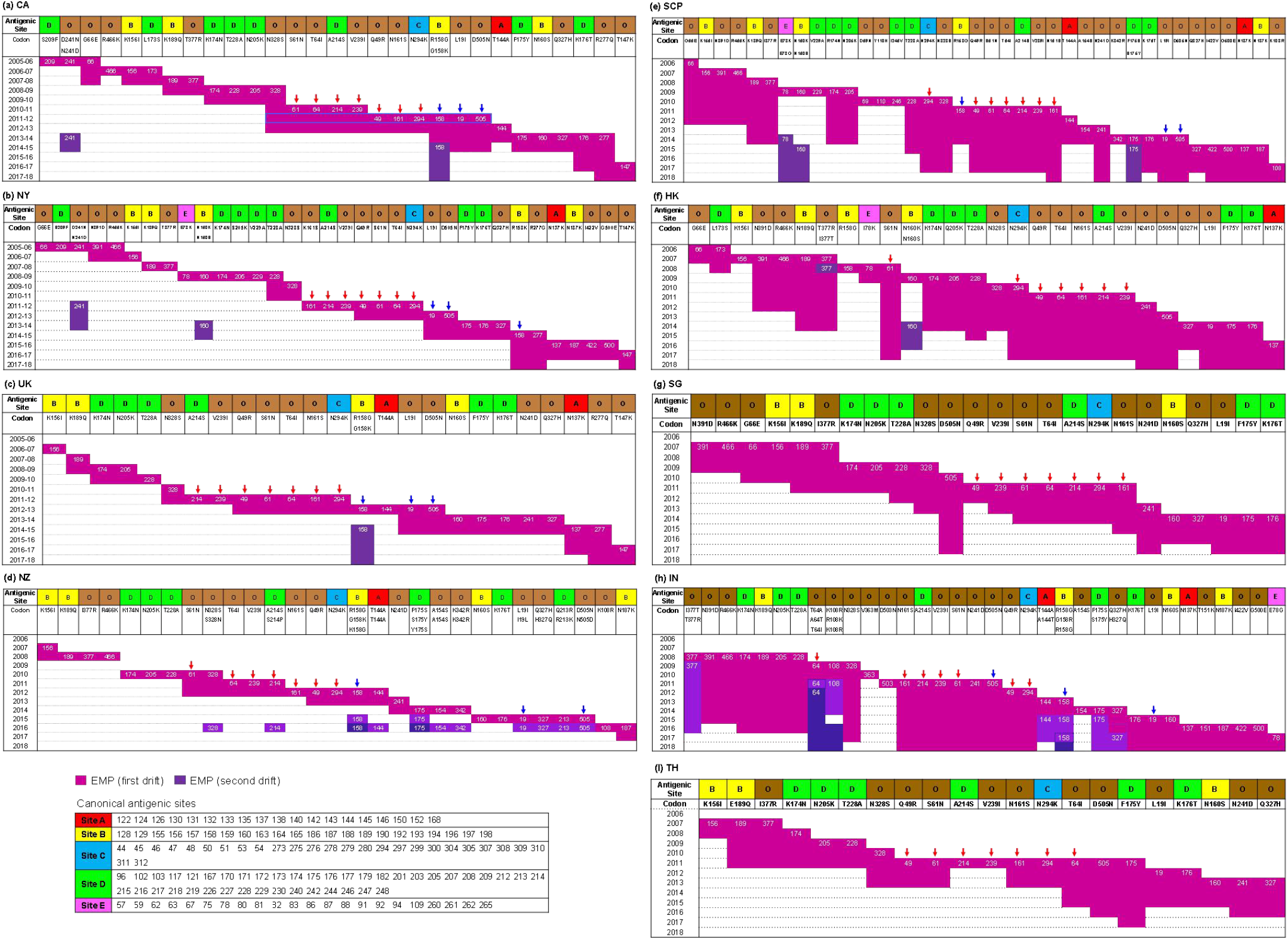
The effective mutations (EMs) in the HA gene of A/H3N2 and their effective mutation periods (EMPs) in six regions from 2005 to 2018. **Legend**: Antigenic sites A–E denoted the canonical antigenic sites at the epitopes. O: others – codon positions not previously reported as antigenic sites. Second row: codon position and subsequent amino acid substitutions, from left to right ranked by occurrence time. The pink and purple colors below the codons indicate the effective mutations (EMs) and their corresponding effective mutation periods (EMP). Red arrow: an example EM set responsible for the 2012–2013 epidemic identified independently in all nine regions. Blue arrow: an example EM set responsible for the 2012–2013 epidemic identified in CA, NY, SCP, UK, NZ and IN.

Overall, the average proportion of EM was only 5.3% of all mutations per year. Among all 46 EMs identified in these 13 years, almost half (43.5%) were from the canonical antigenic sites A–E^3,10,11^ (**Table S1**). Comparing similarity between regions, 93% of the EMs identified in CA were recaptured in NY, while only 41.0% of the EMs were common to all nine regions. The regional unique EMs totaled 15%, contributed mostly by IN and SCP (**Fig. S1**).

### Timing of effective mutations

The genetic epidemiology framework identified a set of advantageous mutations against population immunity during a period of time. The H3N2 is characterized by a continuous global circulation pattern with rapid virus evolution, thus, the timing of mutation event at common EMs could be used to infer the transmission stage of advantageous variant in different regions. In Figure 2, we compared the 15 EMs that were commonly found in at least seven regions and also showed a sequential order of appearance.

**Figure 2.**
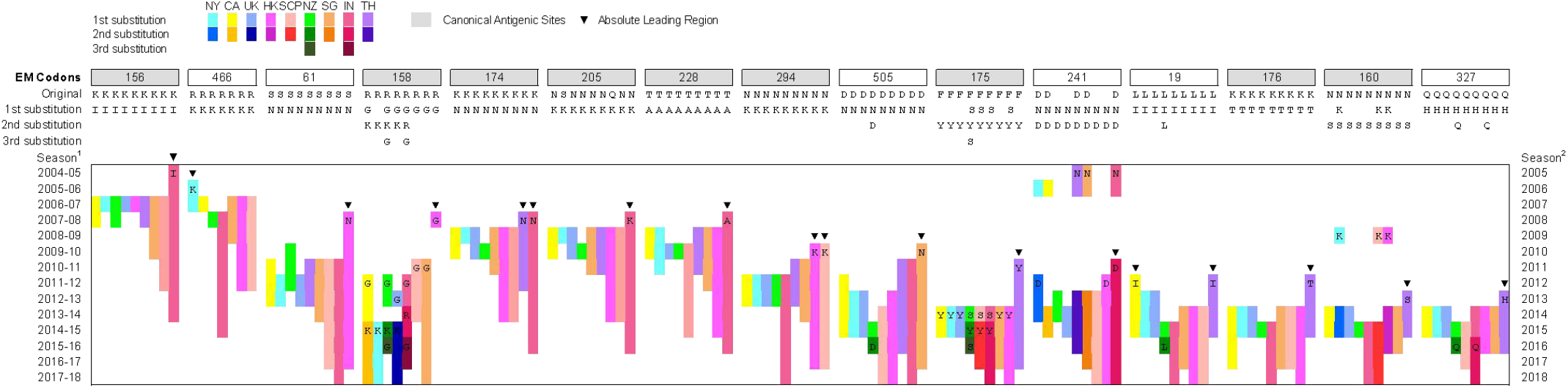
Sequential mutation timing of effective mutations (EMs) in the nine regions. **Legend**: Sequential mutation timing of the EM sites. NY: New York States, CA: California, UK: United Kingdom, HK: Hong Kong SAR, SCP: South China provinces, SG: Singapore, TH: Thailand, IN: India, NZ: New Zealand. Season^1^: influenza season of NY, CA, and UK, from October of one year to April of the subsequent year, e.g., 2005-06 refers to 2005.10 – 2006.04. Season^2^: influenza season of SCP, HK, SG, TH, IN and NZ, span within one year. Details are provided in the Methods and Table S2. 1^st^, 2^nd^, and 3^rd^ substitution: Amino acid substitutions at the same positions, e.g., codon 158 in UK underwent a substitution of R/G in 2012–2013 and was an EM in that year (light blue); the second substitution 158G/K was observed in 2014–2015 and was an EM in 2014–2018 (navy blue).

We found that 90.0% of the common EMs were originated from South East Asia (SEA) (HK, SCP, SG, IN, TH) with an average lead-time of 1.85 years (SD 1.0) relative to North America (NY, CA). While only 10.0% of the EMs were originated from the North America, with an average lead-time of less than one year. According to the sequential appearance order, we could estimate the probability of a region being the source in the viral transmission network, which is the earliest place to observe a mutation, as well as the sink region, where the mutation is lastly observed, and the intersections, which transmission stage lie in between the two. Within SEA, the EMs were most likely to be originated from Thailand (33.3%) and India (30.0%), followed by HK/SCP (16.7%) and SG (3.3%). Previous reports showed that China and SEA lay in the center of the global transmission network^13–16^, and accounted for 66% of the genealogy trunk, estimated using all mutation perturbations on the HA sequences to infer the phylogeny among strains^13^. While we focused on the herd immune evading substitutions that occupy only ~5% of total mutation events on the HA gene. This focused group suggested a stronger role of the SEA region, particularly India and Thailand, in generating the evolutionary advanced genetic variants underlying global H3N2 epidemics. Interestingly, NZ was the most probable place to be the sink region of H3N2 transmission with observed frequency 44.3%, followed by UK (18.1%) and IN (16.9%). Furthermore, NZ and UK were the only two places that had not been a source during the investigation period. While the top connecting sites were SG, NY, CA, SCP, and HK, all regions might act as intersections on the transmission pathways (**Figure 3**). These results provided an explanation to Bedford’s observation that “all regions act to some extent as sources” on the phylogenetic tree. Our results suggested that all regions were intersections, with SEA to be the dominant origin and NZ/UK as the major sink region. The stepwise transmission pathway of advantageous variants was mapped out in Figure 3.

**Figure 3.**
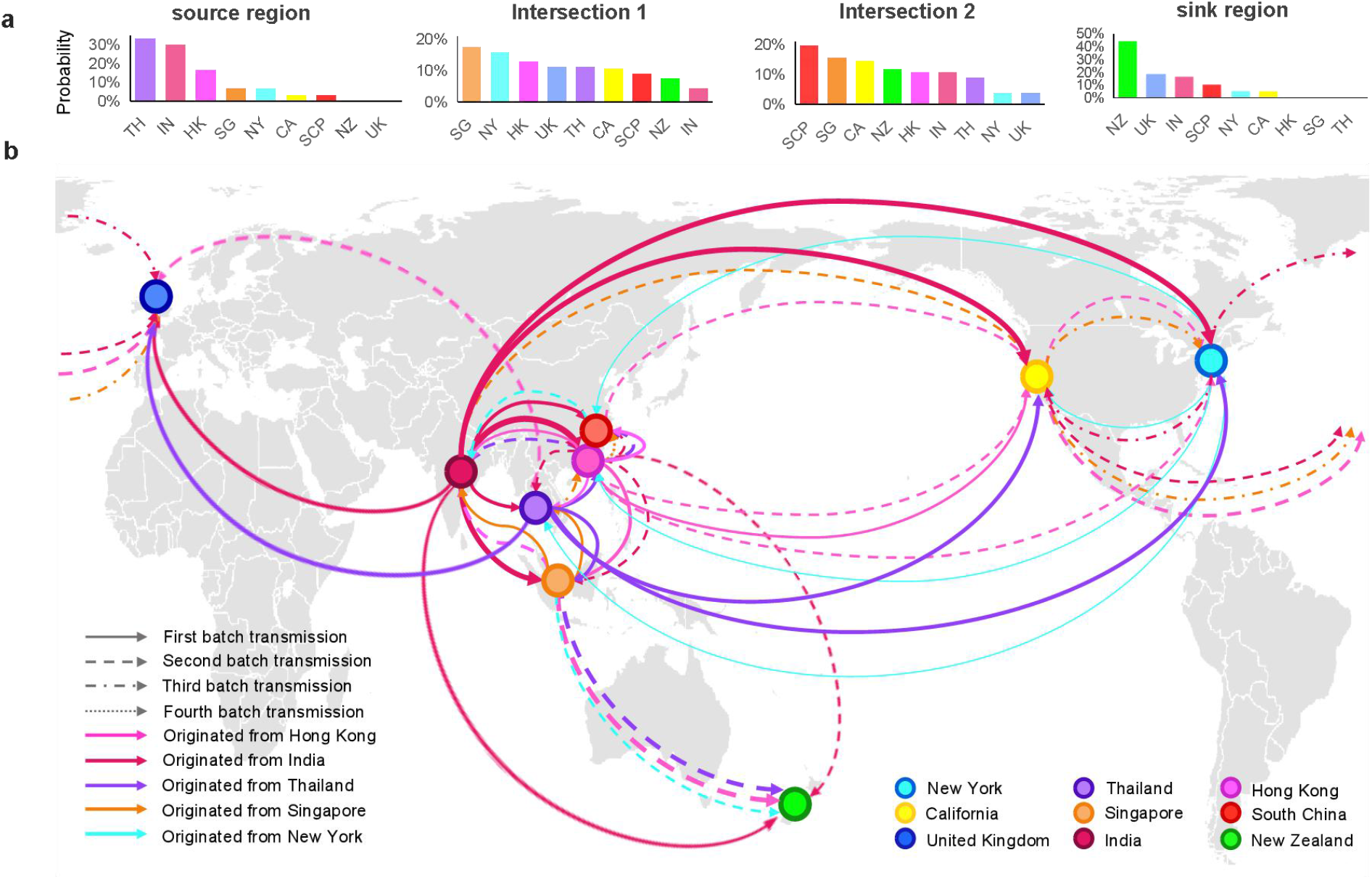
Transmission pathway of influenza virus A/H3N2 inferred by the timing of sequential mutations. **Legend**: NY: New York States, CA: California, UK: United Kingdom, HK: Hong Kong SAR, SCP: South China provinces, SG: Singapore, TH: Thailand, IN: India, NZ: New Zealand. (**a**) Frequency to observe a region in different stages of virus transmission. Overall, for 90% of the times, SEA was observed to be the source region of EMs, while North America was 10%. NZ is the dominant sink region, while all regions might act as connection sites. (**b**) Based on the sequential timing of common EMs, the pathways for viral propagation could be inferred. The line width is proportional to the number of sequential substitutions in the same order.

### The Effective Mutation Period (EMP) – the persistence of viruses with selective advantages

The time duration quantity associated with the EMs, i.e., the EMP, infers the duration of the selective advantage for a new variant in a population, under the influence of herd immunity, migration, and environmental factors. Through the analysis of epidemics in these six regions, we observed distinct characteristics of EMPs from each region. The average EMP estimated in CA, NY, and UK were 2.5 years (SD 1.1), 1.8 years (SD 0.8), and 2.0 years (SD 0.8), respectively. The average EMP determined in the SEA region were considerably longer, being 5.1 years (SD 2.5) for HK, 4.6 years (SD 2.3) for SCP, 4.0 (SD 1.8) for SG, 3.7 (SD 1.5) for TH, and 4.7 (SD 2.9) for IN, thus doubling the length of a set of EMs persisting in NY, CA, and UK. Persistent EMs indicated the long life cycle and persistent circulation patterns of the major strains isolated in SEA region.

## Discussion

In this study, we proposed a novel genetic epidemiology method to identify the effective mutations playing roles in evading population immunity. This is the first method using population epidemic information to infer the statistical importance of amino acids in concurrent viral isolates. In the model, the rising and waning mutation advantage to herd immune response is captured by a site-specific time period, the EMP, providing an unprecedently resolution to mutation period by quantitative method. Since the phenotypic response herein is not the HI titer, the identified amino acids are not confined to those potentially inducing erythrocyte agglutination, and codons contributing to viral propagation, structural stability, and epistasis might be identified. This approach is applicable to thousands of viral strains for selecting important amino acids in any gene segments, subtypes of influenza, and other infectious disease viruses.

Experimentally quantified immune responses displayed a punctuated pattern of evolution, while the genetic evolution, owing to the stochastic nature of mutations, displayed continuous changes^8^. Herein, we distinguished relevant mutations from neutral ones by identifying substitutions contributing to the selective survival advantage of new strains among the human population. Accordingly, the EMs revealed a concordant scale of changes at the epidemic level. The minimum number of EMs observed was 0–2 in the low epidemic season and >10 in the high A/H3N2 epidemic season, with 2.6 EMs per regional epidemic season on average. Smith et al. estimated that 3.6 amino acid substitutions accounted for antigenic drift each year, using combined sequences and worldwide antigenic data^8^. Our estimate is reasonably smaller, as the genetic epidemiological analysis was stratified by the geographical region; therefore, low data heterogeneity resulted in lower genetic variation in regional epidemics. Most EMs were not repetitive during the 13-year study period, indicating an evolutionary tendency of the selection of different amino acid substitutions to generate more antigenically distant strains. In subsequent work, we quantified the importance of the EMs identified here for their roles in stabilizing and shaping phylogenetic tree of HA glycoproteins for the H3N2^17^. A longer observation period in the future would yield additional EMs, enriching the list of key codon positions. Future studies, with increased sampling precision and prolonged data availability, would facilitate further characterization of the evolutionary dynamics of the influenza virus at the molecular level.

## Materials and Methods

### Genetic Data

All A/H3 sequences were retrieved from the National Centre for Biotechnology Information (NCBI)^18^ and the Global Initiative on Sharing all Influenza Data (GISAID)^19^. Sequences longer than 300 codons were retained. The sample size was 863 sequences of New York during 2003-2004 to 2017-2018 flu season, 1,411 of California in 2003-2004 to 2017-2018 flu season, 1,854 of United Kingdom during 2005-2006 to 2017-2018 flu season, 1,628 of SCP during 2003-2018, 849 of Hong Kong during 2005-2018, 1,057 of SG during 2003-2018, 583 of IN during 2003-2018, 849 of TH during 2003-2018 and 733 sequences collected in New Zealand during 2003-2018. The total number of sequences used in the analysis is 9,827. South China provinces include Guangdong, Fujian, Zhejiang, Jiangsu, Anhui, Chongqing, Guangxi, Guizhou, Hainan, Hubei, Hunan, Jiangxi, Shanghai, Sichuan and Yunnan, in which Guangdong and Zhejiang contributed to the majority of samples. The detailed source and sample size of sequence data is in Table S2. The measurements were taken from distinct samples.

### Surveillance Data

The H3 subtype epidemic level is measured by annual sero-positivity rate. The total number of specimens involved in the sero-positivity calculation from governmental reports is 4,104,194^20–28^. Influenza season is defined according to each region’s standard in government report. For New York, California, and United Kingdom, the flu season ranges from October of a year to April next year; for provinces in South China, Hong Kong, Singapore, India and Thailand, the flu season spans the entire year from January to December; and for New Zealand, the flu season ranges from May to September. The detailed source and sample size of surveillance data is in Table S3.

### Weather and population data

For New York, California, United Kingdom, Singapore, India, Thailand and New Zealand, daily relative humidity and temperature are downloaded from public domain^29^. For Hong Kong and South China provinces, relative humidity and temperature are collected from Hong Kong observatory^30^ and China Meteorological Administration^31^, respectively. Meteorological data collection time followed the influenza season defined by the influenza surveillance report. For region including multiple cities, the weather data was collected from the city where largest number surveillance samples were obtained (Table S4 and S5). Population and land area data of the entire region in New York, California, Untied Kingdom, Hong Kong, South China provinces, Singapore, India, Thailand and New Zealand were collected from their government statistical department^32–39^ (Table S6).

### Statistical methods

Below we describe methods that (1) identify the Effective Mutations (EMs) contributing to influenza epidemics; (2) quantify the Effective Mutation Period (EMP), which is the time interval for the effective escape of amino acid substitutions from population immunity. The method involves optimizing two parameters: a dominance threshold *θ* of mutation prevalence; and a time period *h* for mutations to be effective after reaching dominance.

Let *t* denote a time period, such as a year or a month. For each virus sample *i* in period *t*, the observed amino acid sequence is 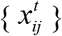, where *j* indicates codon position in the sequence, *j* = 1, …, *J*. Since some codon may have more than one mutations, it would be useful to define a tag sequence to represent all amino acids ever appeared during the investigation period. Let *a_k_* be the tag sequence, *k* = 1,…, *K*. 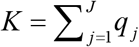, where *q_j_* is the number of unique amino acids observed at codon *j* in the investigation period, and *J* is the total number of codons. Define the coding sequence that translates an observed amino acid sequence to 0/1 format. The coding sequence of a subject *i* at time *t* is 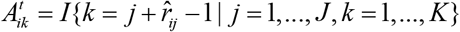, where 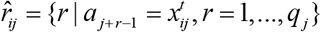. During a time period t, the prevalence of an amino acid at position *k* on the tag sequence is:

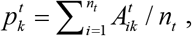

where *n_t_* is the number of sample sequences in *t*. The mutation prevalence vector at *t* is 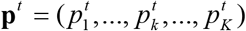.

### Effective mutation and Effective mutation period

The change in the prevalence vector could be used to detect new mutations – a mutation at position *k* can be detected when its prevalence rise from zero at time *t*_0_ to non-zero in a next time *t*_0+1_. While sporadic mutations may appear randomly and do not contribute to virus evolutionary advantage, the effective mutation should show some convincing prevalence in the samples. Therefore, an effective mutation is defined if there exists a time *t*^0^ and a time *t*^θ^, such that the mutation prevalence at codon *k* exceeds a dominance prevalence threshold *θ,* that is, 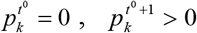, and 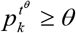, where 0<*t*^0^ ≤*t^θ^*, and *k* =1, …, *K*. The *θ* is estimated by fitting an optimization function matching the mutation activity to population epidemics, 0 ≤ *θ* ≤ 1. When an effective mutation is detected, we can calculate the transition time spent for it to reach dominance, which is 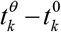. After a mutation reaches dominance, the remaining effective time is denoted as *h*. Therefore, the effective mutation period (EMP) *ω_k_* at a position k includes the transition time and *h*, and is defined as:

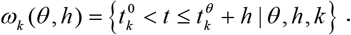

Therefore, the set of effective mutations at time *t* is: 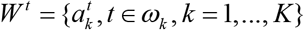. Theoretically, we could estimate a codon specific (*θ*, *h*) estimates if sample size is large enough. Practically, we could reasonably assume that all mutations on a particular gene share the same (*θ*, *h*).

The effective mutations are aggregated to obtain the overall level of genetic activity observed in the sample sequences. Let the effective mutations at time *t* be written as an indicator vector, 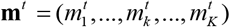, where 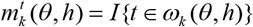. Define the *g*-measure for calculating the overall genetic activity as,

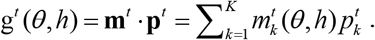

A g-measure vector **g** = [*g^t^*] infers the trend of mutation activity through time, considering the scale and prevalence of mutations.

### Estimating (θ, h)

The (*θ*, *h*) can be estimated by optimizing a fitness function of the g-measure against an epidemic variable, e.g., the sero-positivity rate of a virus subtype. Denote the epidemic response variable as **f** = [*f^t^*]. Let *S*(·) be a function of the fitness between **g** and **f**, e.g., the *p*-value or a goodness-of-fit statistic for a generalized linear model. The optimal estimates for the parameters could be obtained through optimizing,

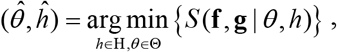

where H={0,1,2, …}, and Θ = [0,1]. Alternative fitting function may be employed when sample size allows. In this study, we used a generalized linear model with sero-positivity rate as response, g-measure as predictor, and the covariates were mean temperature and relative humidity of a region. Figure S2 displayed the parameters fitting heatmap and the optimal parameter combinations. All analysis was conducted in R^40^. All reported *p*-values are two-sided.

## Acknowledgments

This research is supported by NSFC [31871340], Hong Kong HMRF [INF-CUHK-1], and CUHK Direct Grant [4054456] to Maggie H. Wang. We thank Jack Lee, Steven Lau, and Jasmine Choi for their comments on the manuscript.

## Author Contributions

M.H.W designed the study and wrote the manuscript. J.L and L.C did the analysis. M.C.W.C, S.Z, K.C.C, W.K.K.W, R.W.Y.C, B.C.Y.Z, and P.K.S.C commented on the manuscript. W.C and H.Z involved in data collection. E.K.Y approved the paper.

## Additional Information

**Supplement information** is available for this paper in other text.

**Correspondence and requests for materials** should be addressed to M. H. W.

**Reprints and permissions information** is available at www.nature.com/reprints.

