## Supplementary information for "Characterization of the evolutionary dynamics of influenza A H3N2 hemagglutinin"

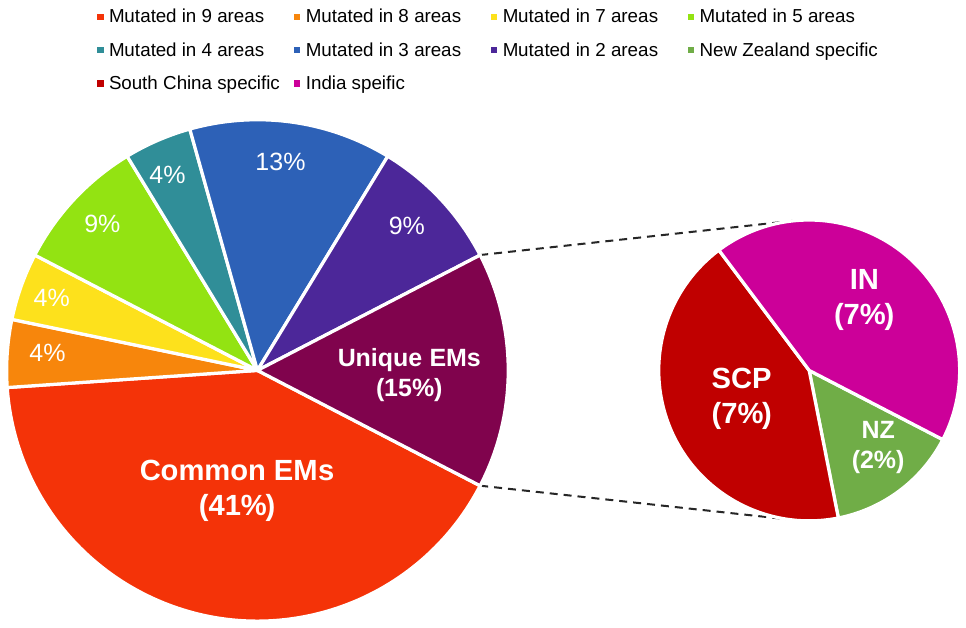

**Fig. S1 | Venn diagram of effective mutations in nine region**s

**Legend:** Among the total 46 EMs identified in nine regions, 41.3% (19/46) were common to all places, 84.8% (39/46) of the EMs was found in at least 2 places, and 15.2% (7/46) EMs were unique to a single place .

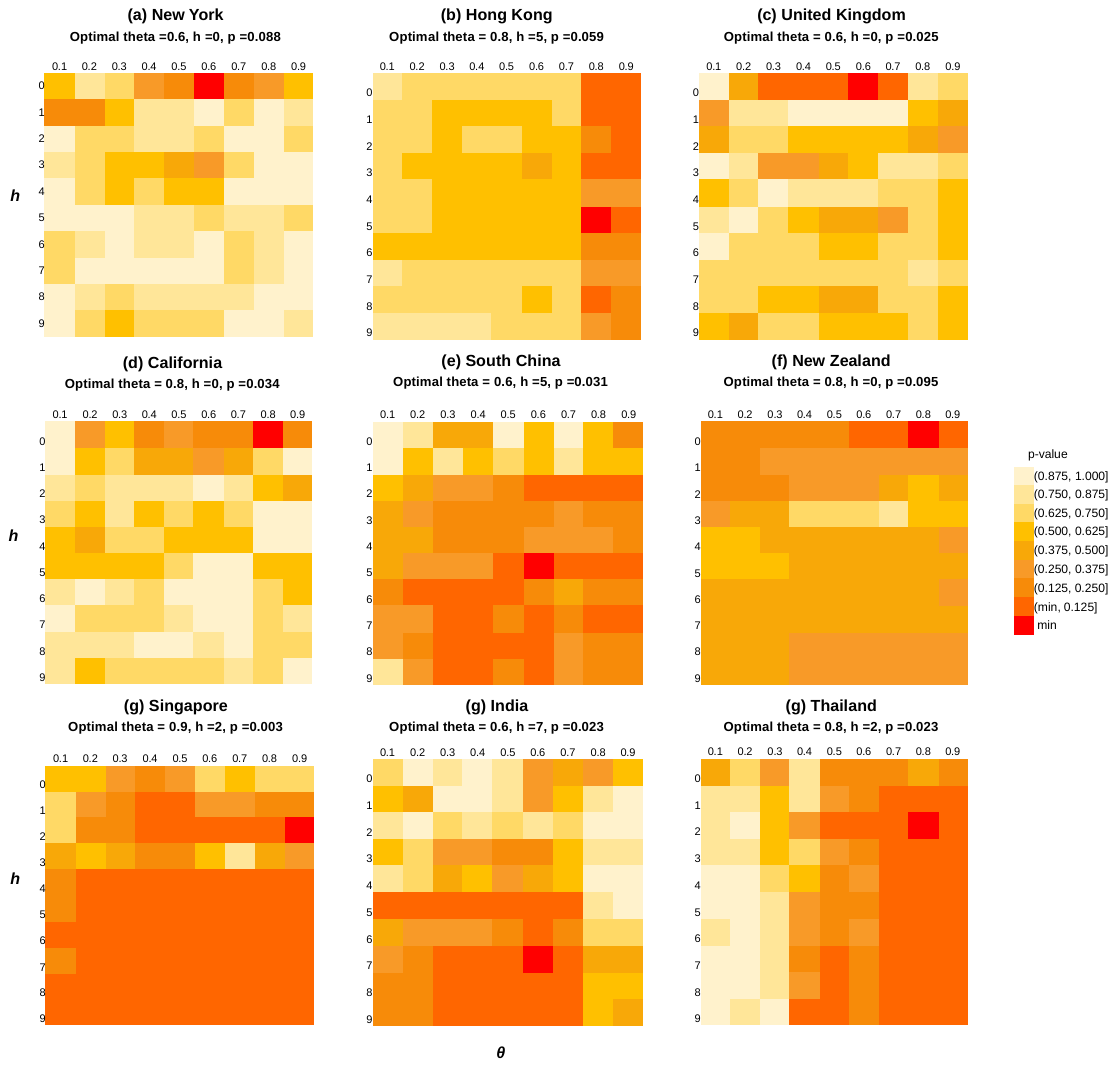

**Fig. S2 | Heatmap of parameter fitting**

**Table S1 |** **List of Effective Mutations**

(A) All EMs:

| **Codon** | **Antigenic site A-E** | **Location of identification** | **HA2** | **Positively selected** |
| --- | --- | --- | --- | --- |
| 19 | - | ALL | - | - |
| 49 | - | ALL | - | - |
| 61 | - | ALL | - | - |
| 64 | - | ALL | - | - |
| 66 | - | NY,CA,HK,SCP,SG | - | - |
| 69 | - | SCP | - | - |
| 78 | E | NY,HK,SCP,IN | - | - |
| 108 | - | SCP,NZ,IN | - | - |
| 110 | - | SCP | - | - |
| 137 | A | NY,UK,HK,SCP,IN | - | Y |
| 144 | A | CA,UK,SCP,NZ,IN | - | - |
| 147 | - | NY,CA,UK | - | - |
| 151 | - | IN | - | - |
| 154 | - | SCP,NZ,IN | - | - |
| 156 | B | NY,CA,UK,HK,NZ,SCP,SG,TH | - | - |
| 158 | B | NY,CA,UK,HK,NZ,SCP,IN | - | - |
| 160 | B | ALL | - | - |
| 161 | - | ALL | - | - |
| 173 | D | CA,HK | - | - |
| 174 | D | ALL | - | - |
| 175 | D | ALL | - | - |
| 176 | D | ALL | - | - |
| 187 | B | NY,SCP,NZ,IN | - | - |
| 189 | B | ALL | - | - |
| 205 | D | ALL | - | - |
| 209 | D | NY,CA | - | - |
| 213 | D | NZ | - | - |
| 214 | D | ALL | - | - |
| 228 | D | ALL | - | Y |
| 229 | D | NY,SCP | - | - |
| 239 | - | ALL | - | - |
| 241 | - | ALL | - | - |
| 246 | D | SCP | - | - |
| 277 | - | NY,CA,UK | - | - |
| 294 | C | ALL | - | - |
| 327 | - | ALL | - | - |
| 328 | - | ALL | - | - |
| 342 | - | SCP,NZ | Y | - |
| 363 | - | IN | Y | - |
| 377 | - | NY,CA,HK,SCP,NZ,SG,IN,TH | Y | - |
| 391 | - | NY,HK,SCP,SG,IN | Y | - |
| 422 | - | NY,SCP,IN | Y | - |
| 466 | - | NY,CA,HK,SCP,NZ,SG,IN | Y | - |
| 500 | - | NY,SCP,IN | Y | - |
| 503 | - | IN | Y | - |
| 505 | - | ALL | Y | - |

NY: New York States, CA: California, UK: United Kingdom, HK: Hong Kong SAR, SCP: South China provinces, NZ: New Zealand, SG: Singapore, IN: India, TH: Thailand.

(B) The nine EMs located on HA2 region: 342, 363, 377, 391, 422, 466, 500, 503 and 505.

**Table S2 | Genetic data sample size - number of sequences**

| **Season** | **NY** | **CA** | **UK** | **Year** | **SCP** | **HK** | **NZ** | **SG** | **IN** | **TH** |
| --- | --- | --- | --- | --- | --- | --- | --- | --- | --- | --- |
| 2002-03 | - | ~~-~~ | ~~-~~ | 2003 | 105 | ~~-~~ | 113 | 36 | 13 | 12 |
| 2003-04 | 87 | 8 | ~~-~~ | 2004 | 165 | ~~-~~ | 125 | 9 | 8 | 16 |
| 2004-05 | 97 | 7 | ~~-~~ | 2005 | 136 | 52 | 79 | 4 | 43 | 25 |
| 2005-06 | 69 | 9 | 12 | 2006 | 24 | 6 | 13 | 5 | 67 | 44 |
| 2006-07 | 30 | 16 | 39 | 2007 | 275 | 7 | 10 | 17 | 40 | 36 |
| 2007-08 | 41 | 28 | 14 | 2008 | 11 | 69 | 12 | 24 | 103 | 33 |
| 2008-09 | 67 | 147 | 13 | 2009 | 18 | 129 | 11 | 39 | 58 | 34 |
| 2009-10 | 11 | 8 | 8 | 2010 | 42 | 53 | 8 | 53 | 17 | 18 |
| 2010-11 | 9 | 19 | 8 | 2011 | 29 | 13 | 16 | 228 | 36 | 76 |
| 2011-12 | 10 | 43 | 47 | 2012 | 49 | 20 | 21 | 7 | 13 | 63 |
| 2012-13 | 48 | 42 | 23 | 2013 | 108 | 29 | 17 | 30 | 68 | 76 |
| 2013-14 | 10 | 39 | 9 | 2014 | 96 | 52 | 15 | 50 | 14 | 85 |
| 2014-15 | 162 | 123 | 411 | 2015 | 170 | 25 | 48 | 65 | 6 | 86 |
| 2015-16 | 47 | 248 | 30 | 2016 | 111 | 22 | 128 | 190 | 5 | 91 |
| 2016-17 | 141 | 347 | 551 | 2017 | 233 | 316 | 104 | 162 | 65 | 91 |
| 2017-18 | 34 | 327 | 689 | 2018 | 56 | 56 | 13 | 138 | 27 | 63 |
| Total | 863 | 1411 | 1854 | Total | 1628 | 849 | 733 | 1057 | 583 | 849 |
| Source | NCBI | GISAID | GISAID | Source | GISAID | GISAID | GISAID | GISAID | GISAID | GISAID |

NCBI: National Center for Biotechnology Information, <https://www.ncbi.nlm.nih.gov/genomes/FLU/Database/nph-select.cgi>

GISAID: Global initiative on sharing all influenza data, <https://www.gisaid.org/>

**Table S3 | Surveillance data sample size - number of specimens**

| **Season** | **NY** | **CA** | **UK** | **Year** | **SCP** | **HK** | **NZ** | **SG** | **IN** | **TH** |
| --- | --- | --- | --- | --- | --- | --- | --- | --- | --- | --- |
| 2002-03 | - | - | ~~-~~ | 2003 | - | - | 8,356 | 1130 | - | - |
| 2003-04 | 16,104 | 467 | - | 2004 | - | - | 789 | - | - | 3910 |
| 2004-05 | 22,534 | 719 | - | 2005 | - | 56,709 | 876 | - | - | 31600 |
| 2005-06 | 20,411 | 876 | 78876 | 2006 | - | 38,391 | 919 | - | - | 5122 |
| 2006-07 | 22,418 | 860 | 75467 | 2007 | - | 56,418 | 772 | 13285 | - | 6200 |
| 2007-08 | 29,495 | 1008 | 59017 | 2008 | 19823 | 43,260 | 947 | 15393 | 3729 | 4811 |
| 2008-09 | 295 | 56204 | 42,482 | 2009 | 97944 | 124,310 | 2,725 | 17037 | 20423 | 6056 |
| 2009-10 | 2,691 | 30773 | 57,198 | 2010 | 84012 | 76,550 | 1,303 | 6926 | 25178 | 7513 |
| 2010-11 | 1,209 | 5718 | 11,249 | 2011 | 42764 | 63,680 | 851 | 2798 | 9833 | 4037 |
| 2011-12 | 702 | 4068 | 31,314 | 2012 | 53660 | 87,590 | 877 | 1526 | 19729 | 4369 |
| 2012-13 | 1,500 | 7754 | 10,680 | 2013 | 109676 | 97,141 | 609 | 1776 | 13119 | 4452 |
| 2013-14 | 2,306 | 11178 | 25,007 | 2014 | 146508 | 110,817 | 733 | 2047 | 7171 | 3733 |
| 2014-15 | 2,463 | 10832 | 32,354 | 2015 | 158262 | 171,249 | 986 | 1926 | 21288 | 1456 |
| 2015-16 | 2,092 | 9391 | 45,366 | 2016 | 152373 | 193,018 | 948 | 2270 | 2151 | 3242 |
| 2016-17 | 1,676 | 10478 | 42,201 | 2017 | 178469 | 225,039 | 2,247 | 2013 | 15091 | 4127 |
| 2017-18 | 2,783 | 12767 | 63,677 | 2018 | 176631 | 255,610 | 1558 | 3379 | 12869 | 3079 |
| Total | 128,679 | 163,093 | 574,888 | Total | 1,220,122 | 1,599,782 | 25,496 | 70,386 | 150,581 | 58,197 |
| Source | NYDOH^i^ | CDPH^ii^   USCDC^iii^ | ECDC^iv^  UKGOV^v^ | Source | CNIC^vi^ | HKCHP^vii^ | NZLMOH^viii^ | GISRS^ix^ | GISRS^ix^ | GISRS^ix^ |

^i^NYDOH: New York Department of Health, <https://www.health.ny.gov/diseases/communicable/influenza/surveillance/>

^ii^CDPH: California Department of Public Health, <https://www.cdph.ca.gov/Programs/CID/DCDC/pages/immunization/flu-reports.aspx>

^iii^USCDC: United States Disease Control and Prevention, <https://gis.cdc.gov/grasp/fluview/fluportaldashboard.html>

^iv^ECDC: European Center for Disease Prevention and Control, <https://ecdc.europa.eu/en/seasonal-influenza>

^v^UKGOV: United Kingdom Government, <https://www.gov.uk/government/collections/weekly-national-flu-reports>

^vi^CNIC: China Influenza Center, <http://ivdc.chinacdc.cn/cnic/>

^vii^HKCHP: Hong Kong Center for Health Protection, <https://www.chp.gov.hk/en/resources/29/304.html>

^viii^NZLMOH: New Zealand Ministry of Health, <https://surv.esr.cri.nz/virology/influenza_surveillance_summary.php>

^ix^GISRS: Global Influenza Surveillance and Response System, <https://www.who.int/influenza/gisrs_laboratory/flunet/en/>

**Table S4 | Meteorological data - Temperature**

| **Season** | **NY** | **CA** | **UK** | **Year** | **SCP** | **HK** | **NZ** | **SG** | **IN** | **TH** |
| --- | --- | --- | --- | --- | --- | --- | --- | --- | --- | --- |
| 2002-03 | - | - | ~~-~~ | 2003 | - | - | 11.15 | 27.81 | - | - |
| 2003-04 | 7.09 | 15.97 | - | 2004 | - | - | 10.72 | 27.89 | - | 29.15 |
| 2004-05 | 7.19 | 15.56 | - | 2005 | - | 23.3 | 11.54 | 28.06 | - | 29.41 |
| 2005-06 | 8.49 | 16.03 | 7.82 | 2006 | - | 23.5 | 10.76 | 27.81 | - | 28.72 |
| 2006-07 | 8.17 | 17.52 | 9.51 | 2007 | - | 23.7 | 11.09 | 27.59 | - | 29.10 |
| 2007-08 | 8.24 | 16.44 | 7.43 | 2008 | 17.53 | 23.1 | 10.78 | 27.54 | 24.50 | 28.28 |
| 2008-09 | 5.95 | 19.54 | 10.89 | 2009 | 17.83 | 23.5 | 11.48 | 27.99 | 25.32 | 28.39 |
| 2009-10 | 9.59 | 16.99 | 6.16 | 2010 | 17.39 | 23.2 | 13.12 | 28.14 | 25.51 | 28.91 |
| 2010-11 | 7.08 | 16.46 | 7.13 | 2011 | 17.21 | 23 | 11.14 | 27.53 | 24.64 | 28.07 |
| 2011-12 | 10.90 | 15.87 | 7.89 | 2012 | 17.11 | 23.4 | 10.58 | 27.62 | 24.81 | 29.64 |
| 2012-13 | 6.67 | 17.00 | 6.28 | 2013 | 18.03 | 23.3 | 11.74 | 27.85 | 24.63 | 29.36 |
| 2013-14 | 7.40 | 17.90 | 8.72 | 2014 | 17.53 | 23.5 | 11.09 | 27.98 | 24.88 | 28.80 |
| 2014-15 | 6.76 | 18.71 | 7.88 | 2015 | 17.48 | 24.2 | 10.68 | 28.29 | 25.21 | 29.21 |
| 2015-16 | 9.80 | 17.86 | 8.40 | 2016 | 18.16 | 23.6 | 11.39 | 28.48 | 25.52 | 29.33 |
| 2016-17 | 8.47 | 17.74 | 8.04 | 2017 | 18.25 | 23.9 | 11.33 | 27.75 | 25.17 | 28.85 |
| 2017-18 | 7.96 | 17.85 | 7.87 | 2018 | 18.12 | 23.9 | 11.14 | 27.79 | 25.11 | 28.75 |
| Average | 7.00 | 17.16 | 8.00 | Average | 17.69 | 23.5 | 11.23 | 27.89 | 25.03 | 28.89 |
| Source | WU | WU | WU | Source | CMD | HKO | WU | WU | WU | WU |

WU: Weather Underground, <https://www.wunderground.com/>

CMD: China Meteorological Administration, <http://www.cma.gov.cn/>

HKO: Hong Kong Observatory, <https://www.hko.gov.hk/contentc.htm>

**Table S5 | Meteorological data - Humidity**

| **Season** | **NY** | **CA** | **UK** | **Year** | **SCP** | **HK** | **NZ** | **SG** | **IN** | **TH** |
| --- | --- | --- | --- | --- | --- | --- | --- | --- | --- | --- |
| 2002-03 | - | - | ~~-~~ | 2003 | - | - | 76.41 | 84.29 | - | - |
| 2003-04 | 65.20 | 66.82 | - | 2004 | - | - | 77.23 | 83.40 | - | 73.85 |
| 2004-05 | 64.55 | 65.89 | - | 2005 | - | 79 | 76.67 | 83.12 | - | 75.61 |
| 2005-06 | 58.74 | 62.73 | 73.40 | 2006 | - | 80 | 77.12 | 84.66 | - | 69.87 |
| 2006-07 | 57.68 | 54.42 | 73.08 | 2007 | - | 77 | 74.13 | 84.56 | - | 67.71 |
| 2007-08 | 60.22 | 57.01 | 73.56 | 2008 | 70.33 | 77 | 79.17 | 83.48 | 66.70 | 67.93 |
| 2008-09 | 59.19 | 58.12 | 74.70 | 2009 | 71.25 | 77 | 76.49 | 82.27 | 61.26 | 66.84 |
| 2009-10 | 62.10 | 58.02 | 81.98 | 2010 | 71.58 | 80 | 78.22 | 83.03 | 65.78 | 67.70 |
| 2010-11 | 59.99 | 59.90 | 77.59 | 2011 | 68.50 | 76 | 76.10 | 84.76 | 67.50 | 70.50 |
| 2011-12 | 60.99 | 60.42 | 78.92 | 2012 | 70.75 | 81 | 74.71 | 84.05 | 61.89 | 69.12 |
| 2012-13 | 61.36 | 59.26 | 78.07 | 2013 | 68.00 | 78 | 79.00 | 82.45 | 68.95 | 67.81 |
| 2013-14 | 56.63 | 56.81 | 77.90 | 2014 | 73.17 | 78 | 77.46 | 79.42 | 65.46 | 70.30 |
| 2014-15 | 56.00 | 58.63 | 75.17 | 2015 | 75.00 | 79 | 76.08 | 77.81 | 66.81 | 69.76 |
| 2015-16 | 56.18 | 56.35 | 76.38 | 2016 | 75.03 | 81 | 76.97 | 76.89 | 66.74 | 72.53 |
| 2016-17 | 59.63 | 61.75 | 75.50 | 2017 | 70.86 | 78 | 79.85 | 83.32 | 66.09 | 74.63 |
| 2017-18 | 61.30 | 56.38 | 76.24 | 2018 | 73.49 | 77 | 78.49 | 80.43 | 66.25 | 75.69 |
| Average | 60.33 | 59.50 | 76.35 | Average | 71.63 | 78.4 | 77.13 | 81.87 | 65.76 | 70.04 |
| Source | WU | WU | WU | Source | CMD | HKO | WU | WU | WU | WU |

WU: Weather Underground, <https://www.wunderground.com/>

CMD: China Meteorological Administration, <http://www.cma.gov.cn/>

HKO: Hong Kong Observatory, <https://www.hko.gov.hk/contentc.htm>

**Table S6 | Population Density**

| **Region** | **Population** | **Land Area(km^2^)** | **Population Density(n/km^2^)** | **Source** |
| --- | --- | --- | --- | --- |
| **NY** | 19,378,102 | 141,300 | 137.14 | USCB - 2010 census |
| **CA** | 37,253,956 | 423,970 | 87.87 | USCB - 2010 census |
| **UK** | 63,181,775 | 243,610 | 259.36 | ONS - 2011 census |
| **SCP** | 768,967,961 | 2,408,514 | 319.27 | NBS - 2010 census |
| **HK** | 7,336,585 | 2,755 | 2663.01 | CSD - 2016 census |
| **NZ** | 4,353,198 | 268,021 | 16.24 | Stats NZ - 2013 census |
| **SG** | 5,076,700 | 721.5 | 7036.31 | SDS - 2010 census |
| **IN** | 1,210,854,977 | 3,287,000 | 368.17 | ORGI - 2011 census |
| **TH** | 65,493,298 | 513,120 | 127.64 | NSO - 2010 census |

UCSB: United States Census Bureau, <http://www.census.gov/>

ONS: Office for National Statistics, <https://www.ons.gov.uk/>

NBS: National Bureau of Statistics, <http://www.stats.gov.cn/english/>

CSD: Census and Statistics Department, <https://www.censtatd.gov.hk/home.html>

Stats NZ: Statistics New Zealand, <https://www.stats.govt.nz/>

SDS: Singapore department of statistics, [www.singstat.gov.sg](http://www.singstat.gov.sg)

ORGI: Office of the Registrar General & Census Commissioner, India, <https://censusindia.gov.in/>

NSO: National Statistical Office, Thailand, <http://popcensus.nso.go.th/en/>
